# Nonlinear averaging of thermal experience predicts population growth rates in a thermally variable environment

**DOI:** 10.1101/247908

**Authors:** Joey R. Bernhardt, Jennifer M. Sunday, Patrick L. Thompson, Mary I. O’Connor

## Abstract

As thermal regimes change worldwide, projections of future population and species persistence often require estimates of how population growth rates depend on temperature. These projections rarely account for how temporal variation in temperature can systematically modify growth rates relative to projections based on constant temperatures. Here, we tested the hypothesis that population growth rates in fluctuating thermal environments differ from growth rates in constant conditions, and that the differing thermal performance curves (TPCs) can be predicted quantitatively. With experimental populations of the green alga *Tetraselmis tetrahele*, we show that nonlinear averaging techniques accurately predicted increased as well as decreased population growth rates in fluctuating thermal regimes relative to constant thermal regimes. We extrapolate from these results to project critical temperatures for population growth and persistence of 89 phytoplankton species in naturally variable thermal environments. These results advance our ability to predict population dynamics in the context of global change.

## Introduction

Organisms live in variable environments. Demographic rates and outcomes that integrate temporal or spatial environmental variation may differ substantially from what might be predicted based on short-term physiological responses to constant, non-varying experimental environments. For example, population growth rates are predicted to vary with temperature as described by their thermal performance curve (TPC), often equated to an organisms’ thermal niche (Figure 1A). Parameters of the thermal niche, such as the upper and lower critical temperatures for population growth, or the optimal temperatures for maximum rates of population growth, are important parameters in the large and growing body of synthesis research that links physiological processes with projected population responses to climate change [1–3]. However, elements of the thermal niche are often documented in physiological assays that use constant laboratory environments. Thermally variable environments can lead to population growth rates over time that differ substantially from estimates based on the average temperature over the same time period [4,5]. This difference complicates projections of population performance based on physiological assays under constant conditions [6], prompting calls for ecologists to explicitly incorporate environmental variation into predictions and models of population and species’ performance in the field [7]. Because temporal patterns of environmental variability differ across regions and the lifespans of organisms, an approach that allows quantitative scaling from thermal performance curves of population growth generated under constant laboratory conditions to population performance in a variable thermal regime may be particularly useful for understanding patterns of abundance and distribution, and species’ responses to climate change.

**Figure 1.**
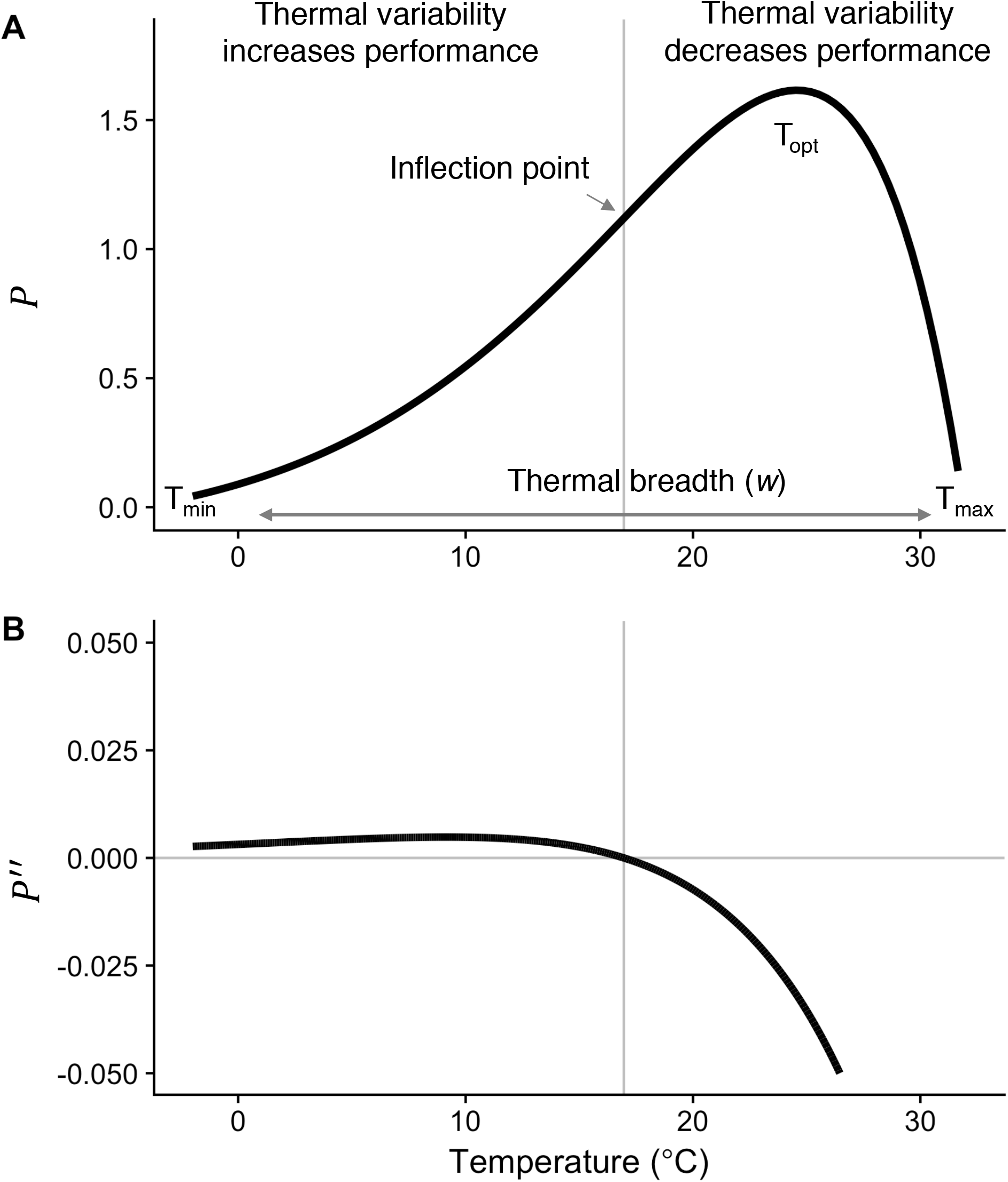
A) A thermal performance curve and its critical temperatures (*T_min_, T_max_, T_opt_*) and thermal breadth, (*w*). The critical temperatures are not fitted parameters of the TPC, but are estimated by numerical optimization. This negatively-skewed curve shows an exponential increase typical of processes following an Arrhenius function, with an accelerating region to the left of the inflection point (grey vertical line), followed by a decelerating region to the right of the inflection point. Notice that the accelerating region corresponds to the region with a positive second derivative (B). Predictions for the temperature ranges in which thermal variability is expected to increase and decrease performance relative to constant conditions are shown. Figure adapted from [7], but parameterized with *T. tetrahele* data from this study.

Biological responses to environmental variation depend on whether the relationship between performance and an environmental gradient is linear or nonlinear [5,8,9], and if nonlinear, whether it is accelerating with increasing temperature, or decelerating (Figure 1A). When performance, *P*, changes nonlinearly with environmental conditions, *E*, time-averaged performance in a variable environment 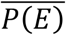 does not necessarily equal performance at the mean environmental condition *P*(*Ē*). This fact, captured by the well-known mathematical rule ‘Jensen’s inequality’, leads to clear predictions about how environmental variability should affect performance over time [8,10]. Jensen’s inequality states that if *P* is a nonlinear function of *E*, then 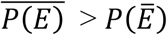 where *P*(*E*) is accelerating (i.e. positive second derivative) and 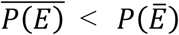 where *P*(*E*) is decelerating (i.e. negative second derivative) (Figure 1B). In the context of temperature, the relationship between organismal or population performance and temperature, captured in the thermal performance curve (Figure 1A), is almost always nonlinear [11] so the often implicitly assumed linear relationship between environment and population growth in demographic models is inadequate to describe population dynamics over temperature gradients [12]. The potential ecological and evolutionary effects of Jensen’s inequality have been shown in several recent studies [1,6,13,14]. Yet, ecologists struggle to incorporate thermal variability when making predictions about the effects of temperature on growth, abundance, and distributions of species in nature, often assuming that species’ thermal experiences are well represented by the mean temperature of their environment.

The typical shapes of TPCs (Figure 1A) [15], with an accelerating phase at lower temperatures and a decelerating phase at higher temperatures within a thermal performance curve suggests positive effects of thermal variation at low temperatures and negative effects at high temperatures [7,14]. Current estimates of the consequences of temporal thermal variability for population-level performance such as population growth rate have assumed a certain shape to the curve (i.e. a Gaussian rise and a parabolic fall; [1]) thus forcing certain outcomes of temporal variability. New evidence suggests that the shape and skew of the TPC can vary substantially among species, phenotypes, or contexts [16], leading to potentially more nuanced responses to environmental variation that may be predicted from empirical thermal performance curves. To date, empirical tests of how temporal temperature variability affects population growth rates have been based on tests at only two temperatures [17]. This restricted sampling of the TPC has precluded testing of quantitative predictions based on the curvature of the TPC.

Here we tested whether population growth in a temporally variable thermal environment reflects the effects of nonlinear averaging of performance at each temperature experienced. For a fast-growing green alga, we tested whether TPCs for population growth generated under constant conditions could predict the outcome of population growth in thermally fluctuating environments. We hypothesized that for populations of this alga, which has overlapping generations and short generation times, population growth measured over several generations would reflect the instantaneous effects of time-averaging of acute thermal responses. Alternatively, if time-dependent stress or acclimation effects that depend on recent thermal history modify growth rates in fluctuating environments [18–20], then population performance in naturally variable environments may not be predicted directly from TPCs generated under constant laboratory conditions, and would require a more detailed understanding of the mechanisms and time-course of thermal niche plasticity.

Following from Jensen’s inequality, we predicted that increased temperature variability over the range of cold temperatures to the left of the inflection point of the TPC (i.e. in the accelerating portion of the curve, where the second derivative of the TPC is positive; Figure 1B) would lead to higher growth rates relative to constant conditions, and that temperature variability in the range of warm temperatures to the right of the inflection point would be detrimental (i.e. where the second derivative of the TPC is negative; Figures 1B and 2A, dashed curve). Then, drawing on a global dataset of empirical TPCs for phytoplankton population growth rates, which vary in shape (skew and position) as well as geographic origin, we use nonlinear averaging (Equation 1) to estimate *in situ* population growth rates, given levels of *in situ* environmental variation at each species’ isolation location. We estimate the extent to which predicted growth rates at each isolation location differ when they are predicted using nonlinear averaging of fluctuating temperatures over time, as compared to when they are predicted based on mean annual temperatures only. By including a range of phytoplankton TPC shapes from a global distribution, we explore the consequences of considering thermal variability in projections of population growth rates, with implications for patterns of abundance and distribution.

**Figure 2.**
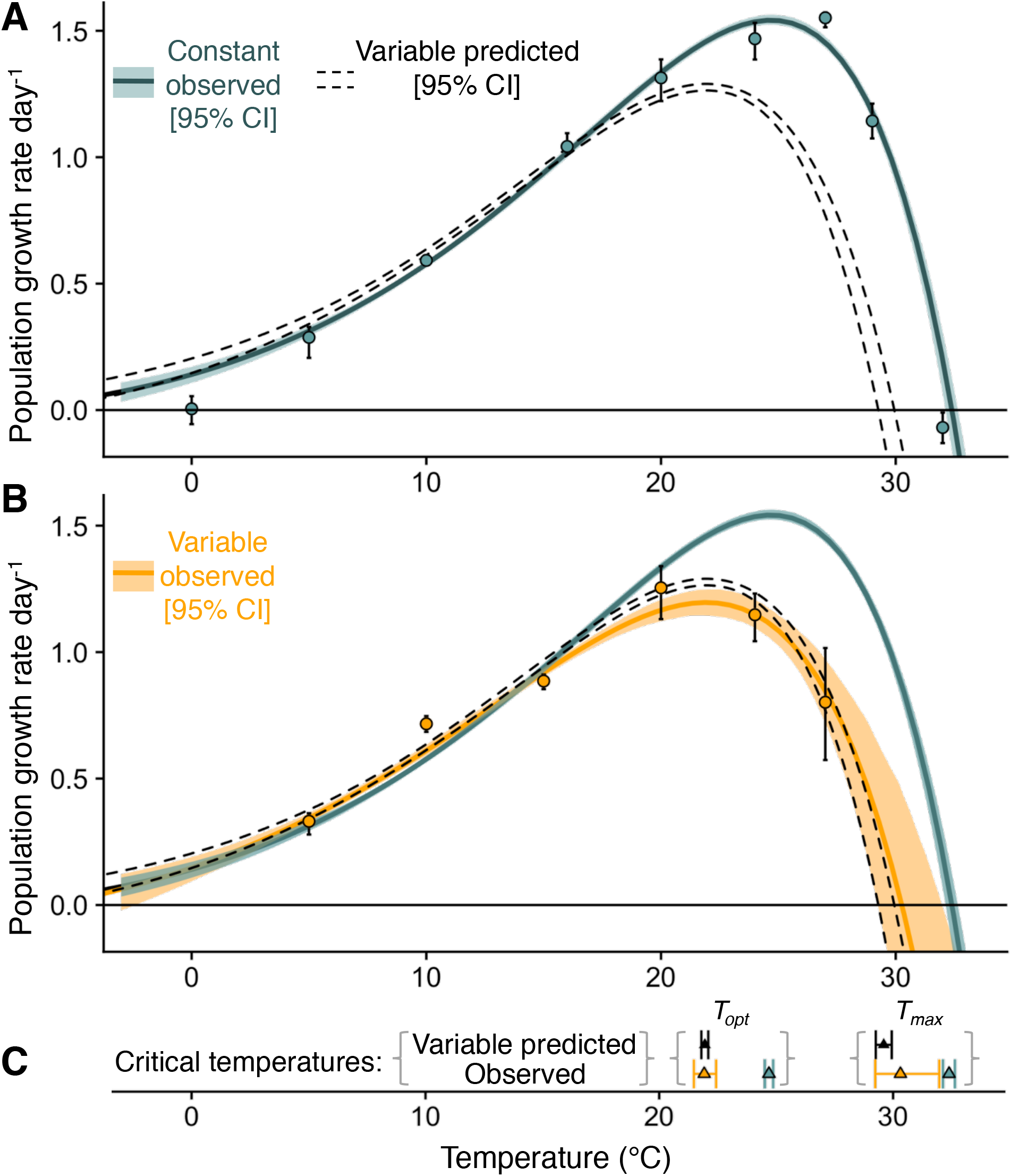
Thermal performance curves for *T. tetrahele* populations growing under constant and variable temperature conditions. A) Exponential growth rates under constant temperature conditions; green line is the fitted thermal performance curve and green shading corresponds to 95% CI generated from non-parametric bootstrapping. The dashed curve represents predicted growth rate under thermally variable conditions based on nonlinear averaging (Equation 1). Points and error bars are observed mean growth rates generated by estimating exponential growth rates at each mean temperature separately (‘indirect’ approach described in the ESM, Appendix A; error bars represent 95% CI). B) Exponential growth rates under thermally fluctuating conditions (±5°C); dashed curve is predicted based on panel A and Equation 1, orange line is the fitted thermal performance curve under fluctuating conditions, orange shading corresponds to 95% CI generated from non-parametric bootstrapping. Orange points and error bars are observed mean growth rates in the fluctuating temperature regime (estimated via the ‘indirect’ approach, Appendix A; error bars represent 95% CI). C) Predicted and observed 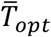 and 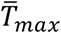 were statistically indistinguishable (predicted: black triangles and 95% CI error bars, observed: orange triangles and 95% CI error bars), and lower than observed *T_max_* and *T_opt_* in constant conditions (green triangles and 95% CI error bars).

## Methods

### Using nonlinear averaging to predict population growth in variable environments

When time series of temperatures experienced by organisms are available and population growth rate, *r* is given by a thermal performance curve, *r* = *f*(*T*), then the expected growth rate, *E*(*r*) averaged over time, *t*, can be calculated by taking the average of the performance at individual time steps:

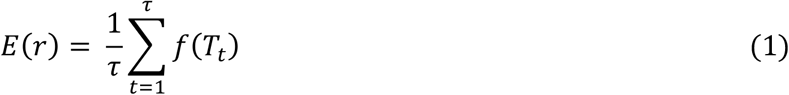

where *t* indexes time.

Empirical records of environmental or body temperatures across time required to use Equation 1 are often not available; however, mean and variance of the distribution of environmental temperatures over a period of time may be more readily accessible. An alternative approach to estimating expected performance under variable conditions when only the mean and variance of the temperature distribution are available is to approximate expected performance under variable conditions using a Taylor approximation of the TPC (Equation S5, an approach that has been incorporated into scale transition theory [7,14,21]). We explore expectations and results using this approach from scale transition theory as well, and compare results using both approaches, to increase the toolkit for ecologists with different kinds of temperature data available (see Electronic Supplementary Material (ESM), Appendix A).

### Experimental quantification of TPCs in constant and fluctuating environments

We experimentally quantified acute TPCs in constant and varying thermal environments for *Tetraselmis tetrahele*, a globally distributed coastal marine phytoplankton species. We used acute TPCs estimated directly from experimental populations, rather than longer-term acclimatized TPCs, to match the time scale of temperature exposure under constant and fluctuating conditions and to simulate the effects of temperature variability in real time. The cultured strain used here was obtained from the Canadian Centre for the Culture of Microorganisms (UW414), and was originally isolated off the coast of Vancouver Island, British Columbia, Canada. The *T. tetrahele* was maintained in laboratory culture in ESAW medium (Enriched Seawater, Artificial Water, [22]) at 16°C on a 16:8 light:dark cycle under nutrient and light saturated conditions before the start of the experiments.

We initiated twenty replicate experimental populations of *T. tetrahele* in 30 mL glass test tubes containing 20 mL of 10uM nitrate ESAW medium at a density of ~ 700 cells/mL under constant temperature conditions at 0°C, 5°C, 10°C, 15°C, 20°C, 24°C, 27°C, 29°C, and 32°C, and under fluctuating temperature conditions with the same mean temperatures as the constant conditions, but fluctuating ± 5°C; i.e. 0°C – 10°C, 5°C – 15°C, 10°C – 20°C, 15°C – 25°C, 19°C – 29°C and 22°C – 32°C. We created fluctuating temperature treatments by programming temperature-controlled incubators (Panasonic MR 154) to switch between the low and high temperature once per day (i.e. approximately 11.5 hours at the high temperature and approximately 11.5 hours at the low temperature, with 30 minutes of transition time in between each temperature cycle). We verified the temperature fluctuations in the experimental populations by measuring water temperatures inside test tubes with 20 mL of water at 1 minute intervals with iButton temperature loggers (Maxim/Dallas Semiconductor)). The fluctuation period of 11.5 hours corresponds to approximately half a generation time at 20°C. To avoid experimental artifacts associated with confounding the temperature cycles with daily light cycles, we grew all experimental populations under 24-hour continuous light, at saturating light intensities of 150 umol/m^2^/s (see Figure S1 in ESM). The source population was acclimated to 24-hour continuous light for four months prior to the experiment. We sampled each of the twenty replicate populations, with four replicates sampled destructively at each of five time points over the period corresponding to the exponential growth phase at each temperature. The intervals between sampling periods depended on the temperature, such that for the warmer temperatures (>15°C) sampling time points were condensed (up to 3 samples per day) to capture exponential growth, while at the colder temperatures (=<15°C) sampling intervals were spread out over longer time periods (i.e. one to five days between sampling points at 0°C and 5°C). Population abundances and cell biovolumes (using area-by-diameter estimation, ABD) were measured from 250 uL samples using a FlowCAM (flow rate = 0.3 ml/min; FlowCAM VS Series, Fluid Imaging Technologies).

### Estimating the temperature dependence of population growth in constant and variable conditions

We estimated the temperature-dependence of population growth directly from the observed time series of population abundance over the temperature gradient [23]. We modeled the temperature-dependent intrinsic rate of population growth, *r*, during the exponential growth phase as:

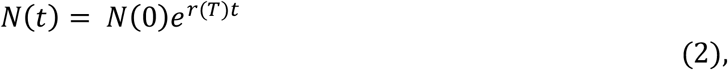

where *N*(*t*) is the number of individuals at time *t*, and *r*(*T*) is given by [16]:

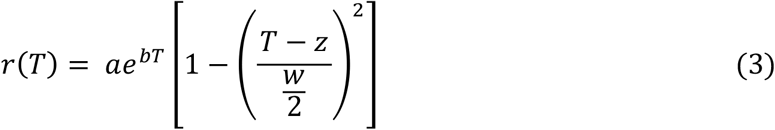

using non-linear least squares regression with the *nls.LM* function in the minpack.LM package in R [24]. Population growth rate, *r*, is a function of temperature, *T, a* and *b* are parameters from the Eppley curve [25] that together describe the increase in maximum observed population growth rates with temperature, *z* determines the location of the maximum of the quadratic portion of the function and *w* is the range over which the growth rate is positive (i.e. the thermal breadth). We also estimated the temperature dependence of population growth via an ‘indirect’ approach [23] (ESM, Appendix A), in which we first estimated growth rates at each temperature separately, and then fit the TPC (Equation 3) to the growth estimates at each temperature. We present the results from this ‘indirect’ approach in the ESM (Figures S2 and S6). To test our hypothesis that performance at constant temperatures predicts performance in fluctuating thermal regimes, we then fit Equations 2 and 3 to the time series of population abundance in the variable experimental treatments, using 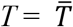, the mean temperature in each treatment in the variable thermal regimes.

Importantly, this generalized TPC equation (Equation 3) does not force the TPC to take on any particular values of *a, b* or *z* or *w*, such that estimated TPCs can be either fully decelerating throughout the entire range of temperatures over which growth is positive, or can have any combination of accelerating and decelerating portions. We favored this TPC equation over others because of this flexibility, because it is parameterized with biologically meaningful parameters, and because has been used previously in studies of phytoplankton TPCs [16,26]. For comparison, we fitted several other functional forms to the experimental dataset (e.g. a Gaussian x Gompertz function, as in [13]) but did not find any better fits when we compared models via AIC.

We quantified four critical temperatures of the TPC (Equation 3) that define the thermal niche (Figure 1A): the optimal temperature for population growth, *T_opt_*, the minimum temperature for positive population growth, *T_min_*, the maximum temperature for positive population growth, *T_max_*, and thermal niche breadth under constant conditions, w. Here we use *T_min_* and *T_max_* to denote the lower and upper limits of the thermal niche for population growth, and note that they are analogous to, but distinct from CTmin and CTmax which are the critical lower and upper limits for organism function [27]. Since *T_opt_, T_min_* and *T_max_* are not parameters of Equation 3, but rather features of the curve, we identified *T_opt_* via numerical optimization using the *optim* function in R and *T_min_* and *T_max_* by finding the roots of the TPC using the *uniroot* function in R. We quantified the analogs of these critical temperatures under thermally variable conditions and refer to them as the minimum mean and maximum mean temperatures for positive population growth under fluctuating conditions, 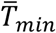 and 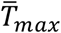 respectively, the mean temperature for optimal growth under fluctuating conditions, 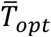, and thermal niche breadth, 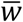. Because *T_min_* from the estimated curve could be below the freezing point of seawater −1.8°C, we used an additional metric of thermal breadth, in which we set *T_min_* to be −1.8°C if it was estimated to be below −1.8°C, because we assumed that *Tetraselmis tetrahele* cannot maintain positive population growth below the freezing point of seawater. We then calculated the range of temperatures over which population growth rate is positive as the difference between *T_min_* and *T_max_*.

To generate estimates of uncertainty in critical temperatures of the TPC (e.g. *T_opt_*) under constant and variable conditions, we determined confidence intervals around fitted thermal performance curves using non-parametric bootstrapping of mean-centered residuals using the *nlsBoot* function with 999 iterations in the *nlstools* package in R. We calculated 95% confidence intervals as the range between the 2.5^th^ and 97.5^th^ quantiles. (Figure 2A, dashed band).

To test our hypothesis that the performance in varying conditions can be explained by nonlinear averaging performance at each temperature experienced, we generated an expected TPC for *T. tetrahele* under thermally fluctuating conditions. We evaluated Equation 1 with *f*(*T*) equal to the TPC fitted using Equation 3, for all values of *T* between 0°C and 33°C (i.e. the entire TPC). We generated confidence intervals around the expected TPC under variable conditions by evaluating Equation 1 for each of the 999 bootstrapped constant-environment curves and calculating 95% confidence intervals as the range between the 2.5^th^ and 97.5^th^ quantiles (Figure 2A, dashed band).

Following Jensen’s inequality, we expected that increased temperature variability should increase population growth in the accelerating phase of the TPC and decrease population growth rate in the decelerating phase of the TPC (Figure 1), and that the shift between positive and negative effects of temperature variability should occur at the inflection point of the constant-temperature TPC. We predicted that increased range of variability should shift the lower and upper limits of the TPC under variable conditions (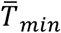 and 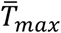) to lower temperatures than *T_min_* and *T_max_* since the TPC is left skewed. We expected the time-averaged maximum growth rate, *r_max_*, to decrease under variable temperature conditions relative to constant conditions because *T_opt_* is always in a decelerating portion of the constant TPC curve. Finally, we expected the thermal breadth under fluctuating conditions, 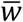, to also decrease under fluctuating conditions if *T_min_* is close to freezing, thus preventing 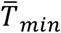 from shifting to lower temperatures to compensate for decreased 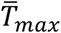.

### *Applying nonlinear averaging to estimate* in situ *phytoplankton population growth rates*

We estimated time-averaged population growth rates in thermally variable environments for a diverse set of phytoplankton species using nonlinear averaging (Equation 1) and scale transition theory (Equation S5). We estimated TPCs for 89 species by fitting Equation 3 to published phytoplankton growth rates [26] measured in the lab at arrays of constant temperatures using maximum likelihood estimation with the *mle2* function in the *bblme* package in R [28] (ESM, Appendix A).

For each of these 89 species, we used historical reconstructed sea surface temperature data to characterize thermal regimes at isolation locations reported in the original studies. For each species’ isolation location, we extracted daily sea surface temperatures (SST) from the closest point in NOAA’s Optimum Interpolation Sea Surface Temperature dataset (OISST), Advanced Very High Resolution Radiometer (AVHRR) and Advanced Microwave Scanning Radiometer on the Earth Observing System (AMSR-E) AVHRR+AMSR, which uses additional data from AMSR-E, available from 2002 to 2011 [29]. This dataset has 0.25° spatial resolution. We calculated the mean and standard deviation of daily sea surface temperatures from 1981 – 2011. We used daily temperature data because these data reflect diurnal and seasonal variation that is central to understanding patterns of long-term population persistence in phytoplankton.

### Statistical estimation of ‘realized’ TPCs

We produced two TPCs for each phytoplankton species – one generated assuming temperatures remain constant through time (akin to constant lab conditions) (‘constant’ scenario), and one that incorporates natural patterns of thermal variability from their habitat (‘variable’ scenario). For the ‘constant’ scenario, we fitted Equation 3 to each species’ population growth rate dataset using the methods described above, and estimated a growth rate, *r*, for the species at the isolation location using *T* = mean annual SST at the isolation location. For the ‘variable’ scenario, we estimated *in situ* growth rates using two approaches: first using Equation 1 where *T_t_* is daily temperature at the isolation location and where *f*(*T*) is the TPC fit using Equation 3, and second using the scale transition theory (Taylor approximation) approach (Equation S5) where *f*(*T*) is the TPC fit using Equation 3, 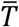 and *σ*^2^_*T*_ are the mean and standard deviation of daily temperatures over the period 1981-2011. Our purpose in using these two approaches was to compare the predictions made with empirical time series of temperature vs. only the mean and standard deviation of the temperature distribution. We present the results from the scale transition theory approach in the ESM. Then, for each species and isolation location, we applied these approaches over the entire TPC, to generate an expected growth rate at each mean temperature assuming a distribution of temperatures that is identical to the daily temperature distribution observed over the historical time period. To do this, we first generated a synthetic temperature distribution around each mean temperature from −2°C to 40°C by taking the distribution of temperatures over the historical time series at each isolation location, subtracting the mean, and then adding each temperature from −2°C to 40°C. We then predicted time averaged growth rates as described above (using Equations 1 and S5). This process yielded a ‘realized’ TPC, which represents expected growth rates given natural patterns of temperature variability. For the ‘constant’ and ‘variable’ scenarios, we compared three attributes of TPCs: *T_opt_, T_min_* and *T_max_*. Given that differences in TPCs based on constant vs variable thermal environments depend on curve shape and temperature variance (Equation S5), we also explored how the discrepancies in estimated critical temperatures and population growth rates at average *in situ* temperatures depend on the shape of the TPC observed in constant lab conditions. We examined the effects of curve attributes including the skew, using a curve skewness metric developed by [26] (Supplementary Equation 5 in [26]), which standardizes the absolute skewness of the curve by the niche width, w, using OLS regression. All analyses were conducted in R (version 3.4.1, [30]). All of the data and code for these analyses are available at https://github.com/JoeyBernhardt/thermal-variability.

## Results

### 1) Does population growth in a thermally variable environment reflect the effects of non-linear averaging over the TPC?

Population growth in fluctuating conditions differed from that in constant thermal conditions over the thermal gradient, and the differences were predicted quantitatively by nonlinear averaging of temporal variation in temperature-dependent performance (95% CI of the growth rate estimates under fluctuating conditions overlapped with predicted growth rates from Equation 1; orange curve and dashed band in Figure 2B). Consistent with expectations based on Jensen’s inequality and nonlinear averaging, experimental populations of *T. tetrahele* had higher population growth rates under fluctuating temperature conditions compared to constant conditions over accelerating portions of the TPC (i.e. at low mean temperatures; 5°C and 10°C), but lower growth rates under fluctuating temperature conditions compared to constant conditions over decelerating portions of the TPC (i.e. at mean temperatures above 10°C, including 15°C, 24°C and 27°C and 29°C) (Figure 2B). Notably, population growth was lower under fluctuating conditions relative to constant conditions at 24°C, which is close to *T_opt_* in this population of *T. tetrahele* (Figure 2B). Populations had negative growth rates at 32°C. The shift between positive effects of temperature fluctuations on population growth at low temperatures and negative effects of fluctuations at warmer temperatures aligned with the inflection point of the constant-temperature TPC (16.76 °C, 95% CI: 16.76°C, 16.81°C), providing strong empirical support for nonlinear averaging in predicting population growth in thermally variable environments.

Thermal variation altered estimated parameter values and key features of the realized thermal performance curve. The maximum exponential growth rate 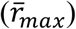 was lower under variable conditions than constant conditions, *r_max_* = 1.54 day^−1^ (95% CI: 1.52 day^−1^, 1.56 day^−1^) vs. 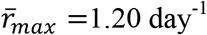 (95% CI: 1.15 day^−1^, 1.25 day^−1^) (Figure 2A, B). Estimated mean optimal temperatures for growth rate were lower under variable conditions: *T_opt_* = 24.69°C (95% CI: 24.52°C, 24.88°C) vs. 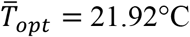 (95% CI 21.48°C, 22.43°C). Maximum mean temperatures for positive growth rates were lower under variable conditions *T_max_* = 32.39°C (95% CI: 32.13°C, 32.64°C) vs. 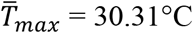 (95% CI: 29.24°C, 31.97°C). All estimated critical temperatures under fluctuating conditions 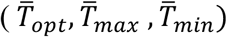 were quantitatively consistent with theoretical predictions from Equation 1 (i.e. had 95% CI overlapping the predicted values from Equation 1; Figure 2B, C). The range of temperatures associated with positive growth rates, accounting for the freezing point of seawater, i.e. *T_max_ − T_min_* was 34.19°C (95% CI: 33.93°C, 34.44°C) under constant conditions and 32.11°C (95% CI: 31.04°C, 33.77°C) under variable conditions. The estimated thermal breadth, w, was also lower under variable conditions, but not statistically distinguishable from constant conditions (i.e. had overlapping 95% CI): 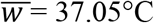, 95% CI: 33.57°C, 45.52°C, vs *w* = 41.23°C, 95% CI: 37.31°C, 47.41°C).

### 2) How different are predicted ‘realized’ TPCs in variable natural environments from predictions based on TPCs generated under constant conditions?

When we estimated the TPCs of the 89 phytoplankton species for constant and varying temperature regimes, we found that for the 90% of species that show negative skew (i.e. mean < median), 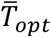 in variable environments is lower than *T_opt_* in constant environments (Figure 3C), while for the remaining 10% of species which show a positive skew, thermal variability is expected to increase 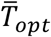 relative to *T*_opt_. The magnitude of the difference between 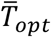 and *T_opt_* increases with increasing standard deviation of sea surface temperature and is well explained by curve skew (slope = 85.98, 95% CI: 70.13, 101.82) and the standard deviation of sea surface temperature (slope = −0.32, 95% CI: −0.40, −0.24) (Adjusted *R*^2^ = 0.66, F_2,86_ = 85.32, *p* < 0.001) (Figure 3C). Phytoplankton growth rate estimates that include the effects of thermal variability, 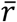, differ from those that do not account for *in situ* thermal variability, *r*, (Figure 3D, F). Generally, predicted growth rates under variable conditions are lower than predicted growth rates assuming constant conditions (i.e. 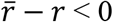, data points below the line y = 0 in Figure 3F), however for some species living in regions with thermal regimes typically colder than the species’ *T*_opt_, growth rates in these environments can exceed those in constant conditions 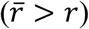. Importantly, the differences between *r*and 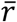 are greatest for species whose isolation locations have mean temperatures that are close to their *T_opt_*. Predicted upper thermal limits for population growth are almost always lower under variable conditions 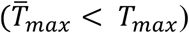 (Figure 3E), and the difference between 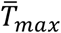 and *T_max_* increases with increasing skewness (positive slope = 53.31, 95% CI: 39.77, 66.85) and standard deviation of SST (negative slope = −0.50, 95% CI: −0.57, −0.43) (Adjusted *R*^2^ = 0.77, *F*_2,75_ = 132.8, *p* < 0.001).

**Figure 3.**
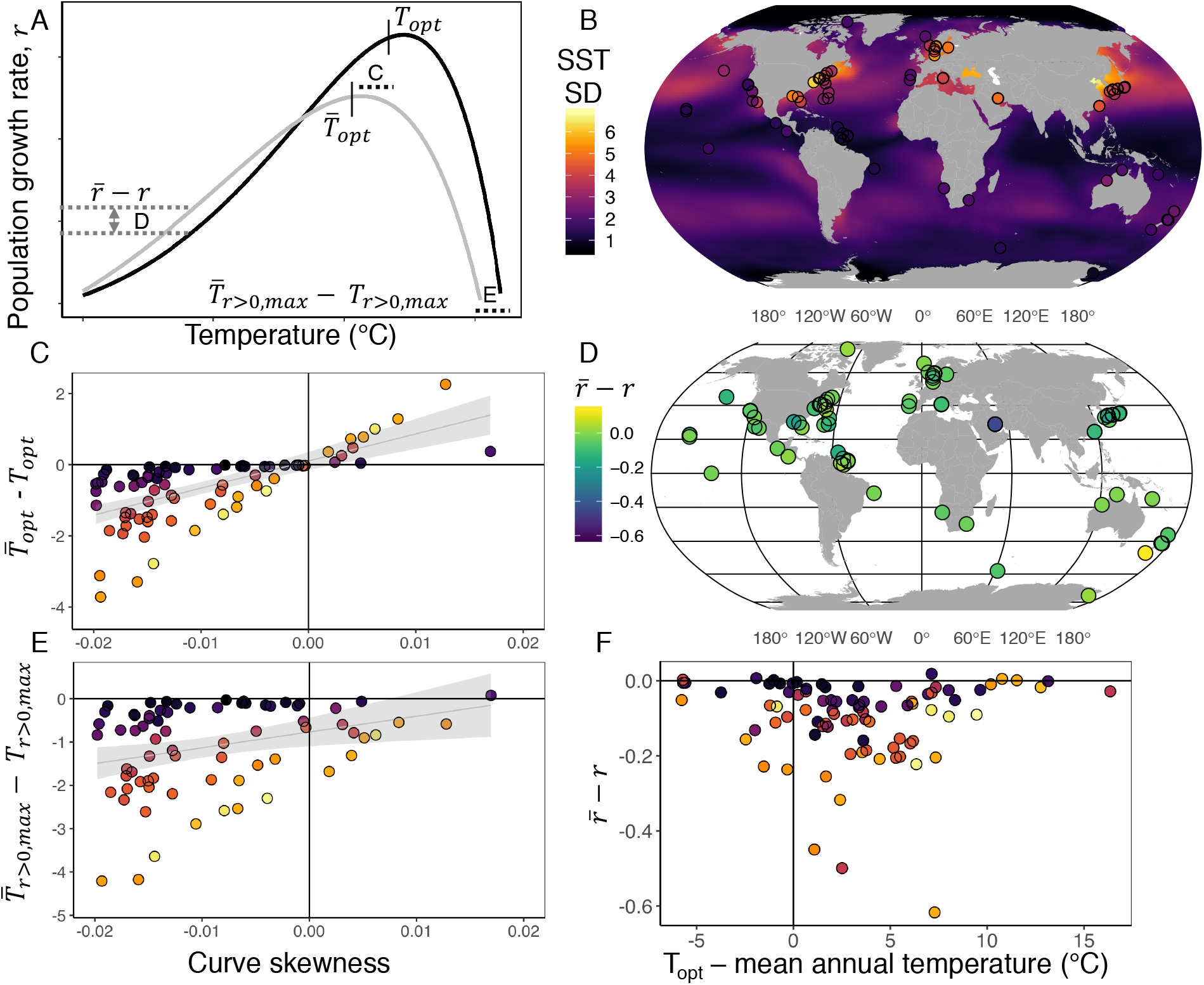
Curve skew and environmental variability predict differences between performance under constant and variable conditions. A) Diagram showing predicted differences in key features of a thermal performance curve under constant (black line) and variable (grey line) environmental conditions. The distances labeled “C”, “D” and “E” illustrate the distances between curve features, plotted in the panels C-E. B) Map of all phytoplankton isolation locations used in the analysis. The color of the ocean and the points corresponds to standard deviation of daily sea surface temperatures over the time period 1981-2011 [29]. C) The difference between predicted 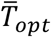 and *T_opt_* generated under constant lab conditions. D and F) Predicted differences in phytoplankton growth rates that do not incorporate *in situ* temperature variation (*r*) vs. predicted growth rates based on Equation 1 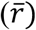. E) The difference between 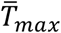 and *T_max_*. Color coding in panels C, E, and F as in B.

For all 89 species in the global dataset, the nonlinear averaging approach (presented here) and the scale transition theory approach (see ESM, Appendix A) resulted in similar ‘realized’ TPCs in variable environments (Figure S4). Predicted critical temperatures, *r_max_* estimates, and relationships shown in Figure 3 were all qualitatively consistent between the two approaches (Figures S4 and S5) indicating that scale transition theory, which makes use of parameters of environmental variation rather than detailed time series, leads to similar predictions in the datasets considered here.

## Discussion

As climate changes worldwide, how temperature affects population growth is a critical link between climate and species persistence in a changing world. One common approach to project population abundance, persistence or fitness under future climate conditions is to apply mathematical curves describing population growth rate over a range of temperatures (a TPC) generated from controlled lab studies (e.g. [31]). This approach relies on the assumption that TPCs do not vary systematically with thermal variation. Here we tested this important assumption and found that natural levels of environmental variability systematically change how population growth depends on temperature. In our analysis of globally distributed phytoplankton TPCs, we found that a variable thermal environment reduced critical upper mean temperatures (*T_max_*) for population persistence by up to 4°C, meaning that population growth in variable conditions was much lower at warmer temperatures than would be predicted based on a TPC generated under constant conditions. This thermal differential is substantial – the 4°C difference in *T_max_* is on par with the magnitude of predicted temperature changes over the next 100 years [32], suggesting that projections of TPCs used for future conditions may overestimate population performance in warming climates. Other work has compared acute thermal physiological limits (e.g. CT_max_, CT_min_) to environmental temperatures at range limits to assess relative sensitivities of range edges to warming (e.g. [3]) yet the underlying nonlinear negatively-skewed thermal performance curve expected for these ectotherms suggests that variability at warm range edges will have a stronger effect on population persistence than variability at cold range edges. Specifically, our findings suggest that approaches based on direct applications of lab-determined critical temperatures may under-predict range edges at boundaries defined by cold temperatures, and over-predict range edges at boundaries defined by warm temperatures.

We have shown experimentally that realized TPCs in variable environments differ from those in constant environments, and that these differences are predicted qualitatively by Jensen’s inequality [8,14] and quantitatively from nonlinear time averaging of performance over the TPC [21]. Fluctuating temperatures changed several aspects of the ‘realized’ thermal performance curve, including the mean temperature of optimal population growth 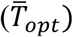 and the maximum growth rate 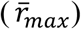 – effectively shifting the TPC toward lower temperatures and lower population growth rates overall. Consistent with the argument that ‘suboptimal’ is optimal [13], we show both experimentally (Figure 2B) and theoretically (using empirical TPCs and *in situ* temperatures; Figure 3D, F) that population growth rates are often lower under variable thermal conditions relative to constant ones, and this negative effect of temperature variation is greatest for populations living close to their thermal optima. However, in contrast to the common assumption that environmental variation is always detrimental for population growth rates [12], our results suggest that populations living at mean temperatures in an accelerating part of the TPC will benefit from environmental variation. Indeed, the *T. tetrahele* isolate used here was collected at a location where mean annual temperatures are far colder than its 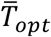, in the accelerating portion of the negatively skewed TPC (at mean temperature = 6.92°C [29]). In this way, when TPCs have accelerating portions at the edges of the thermal niche, thermal variation may allow population persistence in environments that would be too hot or cold under constant conditions.

When we applied nonlinear averaging to estimate the growth rates of globally distributed phytoplankton species, we found that the effect of variability of predicted phytoplankton thermal performance depended strongly on the shape and skew of the TPC and the degree of thermal variability in the oceans from which the phytoplankton originated. Previous approaches have, in the absence of more complete datasets, assumed a certain shape to the reaction norm or TPC (i.e. a Gaussian rise and a parabolic fall; [1]), thus forcing certain outcomes of variability. Here we used a model that does not prescribe any particular shape (i.e. allows for fully decelerating curves, or curves with accelerating portions, or any combination of decelerating and accelerating portions), thus enabling a more complete exploration of the potential effects of temperature variability on population performance. Importantly, empirical TPCs varied in skew, and whether the TPC was positively- or negatively-skewed determined the direction of the shift between *T_opt_* and 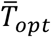. The majority of the curves in the dataset were negatively skewed, and in these cases variability shifted *Ŧ_opt_* to colder temperatures. Negatively skewed TPCs are widely observed across ectothermic taxa [15], suggesting that the direction of the effects of thermal variability observed in our experiment may be general across many ectothermic taxa and ecosystems. Because the shape and skew of the TPC determine performance in variable environments, the mechanisms that determine the shape of the thermal performance curve can have an important influence on the outcome of thermal variability on population persistence. More studies of the diversity of TPC shapes among species will elucidate the extent to which environmental variability increases or decreases performance optima relative to constant lab conditions.

The mathematical tools we applied are generalizable to assessments and projections of biological responses to environmental change and should replace the direct application of TPC parameters based on constant conditions in the lab. In the absence of empirical temperature time series, predictions made based on the mean and standard deviation of the temperature distribution may provide a sufficiently accurate approach. We predicted similar effects of thermal variability on population growth rates when these predictions were made using empirical time series of *in situ* SST (Equation 1) and when using a Taylor approximation approach from scale transition theory (See ESM, Equation S5, and Figures S4-5). Indeed, most of the phytoplankton strains we studied were isolated at locations with thermal regimes corresponding to portions of their TPCs lower in temperature than the highly non-linear temperature ranges near *T_opt_*.

Our results, that performance in fluctuating environments can be predicted from TPCs generated in constant conditions, differ from two previous attempts to predict individual somatic growth rates in fluctuating environments based on TPCs generated in constant conditions [18,19]. Previous observations showed that short-term acute responses to diurnal temperature variation were not predictable based on TPCs generated from chronic exposure to constant temperatures over the course of development of an insect [19] and amphibian [18]. These contrasting results highlight the importance of the time scale of temperature exposures used to measure and predict performance in constant and fluctuating conditions. Biological responses to temperature, including acclimation and thermal stress are inherently time-dependent and may accrue over the course of development in longer lived species [11,33–35], thus precluding the ability of TPCs generated over longer terms (i.e. entire lifespans of individuals) to predict temperature responses over relatively short time spans (small fractions of lifespans corresponding to daily temperature variation). In our experiments, using a fast-growing phytoplankton with short overlapping generations, we maintained the time frames of temperature exposure comparable under both constant and fluctuating conditions. We used *acute* thermal performance curves for population growth rates, which integrate responses to temperature over several generations, generated with a very short acclimation duration at each test temperature, to predict *acute* responses to thermal variability, also over several generations, thus keeping the time frames comparable. By measuring population growth over several generations in both constant and fluctuating conditions, this approach allowed us to avoid mismatches in time scale and time-dependent effects and instead test the nonlinear effects of temperature variation.

The predictions we made of *in situ* phytoplankton population growth rates should be interpreted as first-order predictions, which do not incorporate long-term phenotypic responses to thermal variability. Organisms may be able to acclimatize or adapt to fluctuating conditions over longer-term exposures [36], with the potential to alter the shape and limits of the TPC during the time-course of environmental variability. The global predictions we make here should be viewed as null models which do not incorporate long-term biological responses to environmental variability, and should be tested empirically [6,7]. To extend our predictions of time-averaged growth rates at the isolation location temperatures to time-averaged growth rates over the whole thermal niche, i.e. our visualization of a ‘realized TPC’, we had to assume a particular distribution of temperatures around each hypothetical mean, and used the variation observed at each isolation location as the residual variation around each putative mean temperature. This assumption about temperature distributions is a simplification of real thermal regimes, which likely show more complex patterns of variability and temporal autocorrelation, which can further modify the effects of variability on populations [4]. Resource supply is also likely to covary with temperature, potentially altering the outcomes of thermal variability on population growth rate. Nevertheless, even in the simplest scenario of environmental variability, we predict significant changes in realized mean TPC critical temperatures.

Understanding population responses to temperature now and into the future involves understanding biological responses to changes in the full cassette of temperatures experienced – i.e. all the variation. Omitting the effects of environmental variation from population and species distribution models may limit our ability to predict species’ responses, particularly at the extreme edges of their ranges, even if variability patterns remain unchanged. We show that the effects of environmental variation can be predicted based on the shape of the functional relationship between population growth and the environment, adding another tool to the kit for forecasting species’ responses to the environment in a changing world.

## Acknowledgements

We thank M. Thomas for sharing code used for thermal performance curve fitting, and C. Harley and A. Gehman for helpful comments on previous versions of the manuscript.

**Author contributions:** JRB conceived the study and designed the experiments, with help from JMS, PLT and MIO. JRB carried out the experiments, analyzed the data and wrote the first draft of the manuscript. All authors contributed to writing the manuscript. All authors gave final approval for publication.

## References

1. Vasseur DA, DeLong JP, Gilbert B, Greig HS, Harley CDG, McCann KS, Savage V, Tunney TD, O’Connor MI. 2014 Increased temperature variation poses a greater risk to species than climate warming. Proc. R. Soc. London B Biol. Sci. 281.

2. Buckley LB, Nufio CR, Kingsolver JG. 2014 Phenotypic clines, energy balances and ecological responses to climate change. J. Anim. Ecol. 83, 41–50. (doi: 10.1111/1365-2656.12083)

3. Sunday JM, Bates AE, Dulvy NK. 2012 Thermal tolerance and the global redistribution of animals. Nat. Clim. Chang. 2, 686.

4. Gonzalez A, Holt RD. 2002 The inflationary effects of environmental fluctuations in source-sink systems. Proc. Natl. Acad. Sci. U. S. A. 99, 14872–7. (doi: 10.1073/pnas.232589299)

5. Lawson CR, Vindenes Y, Bailey L, Pol M. 2015 Environmental variation and population responses to global change. Ecol. Lett. 18, 724–736.

6. Estay SA, Lima M, Bozinovic F. 2014 The role of temperature variability on insect performance and population dynamics in a warming world. Oikos 123, 131–140. (doi:10.1111/j.1600-0706.2013.00607.x)

7. Dowd WW, King FA, Denny MW. 2015 Thermal variation, thermal extremes and the physiological performance of individuals. J. Exp. Biol. 218.

8. Ruel JJ, Ayres MP. 1999 Jensen’s inequality predicts effects of environmental variation. Trends Ecol. Evol. 14, 361–366. (doi:10.1016/S0169-5347(99)01664-X)

9. Drake JM. 2005 Population effects of increased climate variation. Proc. R. Soc. London B Biol. Sci. 272.

10. Jensen J. 1906 Sur les fonctions convexes et les inegalites entre les valeurs moyennes.

11. Schulte PM, Healy TM, Fangue NA. 2011 Thermal Performance Curves, Phenotypic Plasticity, and the Time Scales of Temperature Exposure. Integr. Comp. Biol. 51, 691–702. (doi: 10.1093/icb/icr097)

12. Lande R, Engen S, Saether B-E. 2003 Stochastic population dynamics in ecology and conservation. (doi:10.1093/ISBN)

13. Martin TL, Huey RB. 2008 Why suboptimal is optimal: Jensen’s inequality and ectotherm thermal preferences. Am. Nat. 171, E102–18. (doi:10.1086/527502)

14. Denny M. 2017 The fallacy of the average: on the ubiquity, utility and continuing novelty of Jensen’s inequality. J. Exp. Biol. 220, 139–146. (doi:10.1242/jeb.140368)

15. Dell AI, Pawar S, Savage VM. 2011 Systematic variation in the temperature dependence of physiological and ecological traits. Proc. Natl. Acad. Sci. 108, 10591–10596.

16. Thomas MK, Kremer CT, Klausmeier CA, Litchman E. 2012 A global pattern of thermal adaptation in marine phytoplankton. Science (80-.). 338, 1085–1088.

17. Bozinovic F, Bastías DA, Boher F, Clavijo-Baquet S, Estay SA, Angilletta MJ. 2011 The mean and variance of environmental temperature interact to determine physiological tolerance and fitness. Physiol. Biochem. Zool. 84, 543–52. (doi:10.1086/662551)

18. Niehaus AC, Angilletta MJ, Sears MW, Franklin CE, Wilson RS. 2012 Predicting the physiological performance of ectotherms in fluctuating thermal environments. J. Exp. Biol. 215, 694–701. (doi:10.1242/jeb.058032)

19. Kingsolver JG, Higgins JK, Augustine KE. 2015 Fluctuating temperatures and ectotherm growth: distinguishing non-linear and time-dependent effects. J. Exp. Biol. 218, 2218–25. (doi:10.1242/jeb.120733)

20. Kremer CT, Fey SB, Arellano AA, Vasseur DA. 2018 Gradual plasticity alters population dynamics in variable environments: thermal acclimation in the green alga *Chlamydomonas reinhartdii.* Proc. R. Soc. B Biol. Sci. 285, 20171942. (doi:10.1098/rspb.2017.1942)

21. Chesson P, Donahue MJ, Melbourne B, Sears ALW. 2005 Scale Transition Theory for Understanding Mechanisms in Metacommunities. In Metacommunities: spatial dynamics and ecological communities, pp. 279–306.

22. Harrison PJ, Waters RE, Taylor FJR. 1980 A broad spectrum artificial sea water medium for coastal and open ocean phytoplankton. J. Phycol. 16, 28–35.

23. Palamara GM, Childs DZ, Clements CF, Petchey OL, Plebani M, Smith MJ. 2014 Inferring the temperature dependence of population parameters: the effects of experimental design and inference algorithm. Ecol. Evol. 4, 4736–4750. (doi:10.1002/ece3.1309)

24. Elzhov T V, Mullen KM, Spiess A-N, Maintainer BB. 2016 Title R Interface to the Levenberg-Marquardt Nonlinear Least-Squares Algorithm Found in MINPACK, Plus Support for Bounds.

25. Eppley RW. 1972 Temperature and phytoplankton growth in the sea. Fish. Bull 70, 1063–1085.

26. Thomas MK, Kremer CT, Litchman E. 2016 Environment and evolutionary history determine the global biogeography of phytoplankton temperature traits. Glob. Ecol. Biogeogr. 25, 75–86. (doi:10.1111/geb.12387)

27. Lutterschmidt WI, Hutchison VH. 1997 The critical thermal maximum: history and critique. Can. J. Zool. 75, 1561–1574. (doi:10.1139/z97-783)

28. Bolker B. 2017 bbmle: Tools for General Maximum Likelihood Estimation.

29. Reynolds RW et al. 2007 Daily High-Resolution-Blended Analyses for Sea Surface Temperature. J. Clim. 20, 5473–5496. (doi:10.1175/2007JCLI1824.1)

30. R Core Team. 2017 R: A Language and Environment for Statistical Computing.

31. Deutsch CA, Tewksbury JJ, Huey RB, Sheldon KS, Ghalambor CK, Haak DC, Martin PR. 2008 Impacts of climate warming on terrestrial ectotherms across latitude. Proc. Natl. Acad. Sci. 105, 6668–6672.

32. Frölicher TL, Paynter DJ. 2015 Extending the relationship between global warming and cumulative carbon emissions to multi-millennial timescales. Environ. Res. Lett. 10, 75002.

33. Kingsolver JG, Woods HA. 2016 Beyond Thermal Performance Curves: Modeling Time-Dependent Effects of Thermal Stress on Ectotherm Growth Rates. Am. Nat. 187, 283–294. (doi:10.1086/684786)

34. Kingsolver JG, Buckley LB. 2015 Climate variability slows evolutionary responses of Colias butterflies to recent climate change. Proceedings. Biol. Sci. 282. (doi:10.1098/rspb.2014.2470)

35. Kingsolver JG, Buckley LB. 2017 Quantifying thermal extremes and biological variation to predict evolutionary responses to changing climate. Philos. Trans. R. Soc. Lond. B. Biol. Sci. 372, 20160147. (doi:10.1098/rstb.2016.0147)

36. Angilletta MJ. 2009 Thermal Adaptation: A Theoretical and Empirical Synthesis. (doi: 10.1093/acprof:oso/9780198570875.001.1)

